# Influence of Organic Matrix and Cations on Bio-Methane Yield with Anthracite Methanogenic Consortium

**DOI:** 10.1101/555532

**Authors:** Dong Xiao, Enyuan Wang, Yidong Zhang

## Abstract

Organic compounds fermentation of coal has been used to generate secondary biogenic gas and enhance gas reservoirs in coalbed. To enhance the bio-degradation process, culture nutrition plays an important role in remediating the nutritional deficiency of the coal seam. The influence of bio-methane yield with organic inputs and cations concentrations was examined. Research of organic matrix influence revealed that the traditional organic material except yeast extract should forbid, and the input of yeast extract should limit at 1.00g/L also. Further, the study demonstrated that the ion concentration of sodium, potassium, magnesium, calcium and ammonia nitrogen also influenced methane and carbon dioxide yields. And the optimize concentrations for Ca^2+^, K^+^, Na^+^, Mg^2+^ were 5.1, 1.7, 23 and 1.3 mmol/L. The Mg^2+^ was particularly sensitive in inhibiting CH_4_ metabolism processes largely for gas-coal methanogenic consortium.

## 1. INTRODUCTION

The demand for gas has seen a tremendous increase in the last 10 years [1]. In particular, coal-bed methane enhance with biotechnology represents an ideal method to increase methane productivity from oil deposits and coal seams using underground anaerobic digestion processing [2,3]. With the addition of adequate nutrient supplements, methanogenic consortium has the potential to degrade organic compounds in coal into methane. In this way, gas reserves and productivity could be enhanced along with secondary biogenic gas generation [4–6].

Previous studies involved with the methanogenic microbial traits and fermentation characteristics for bio-methane generation have focused on the processes of bio-degradation by microbial communities in coal seam [2,7,8] Anion research identified that methane biosynthesis process could active where coalbed waters exhibit relatively low salinities (<2 mol/L Cl^−^) and low SO_4_^2−^ concentrations (<10 mmol/L) [9]. Nutrients supply by meteoric water plays an important role in methanogenic microbial metabolism [10,11]. However, nutritional deficiency exists in most coal seam. Most organic compounds in coal cannot be readily used as nutrients for methanogenic microbes. In addition to the inherent elements present in coal beds, supplemental nutrients are important to remediate the nutritional deficiency and promote microbial growth [12,13]. If concentrations of supplemental nutrients are excessive, the methanogenic consortium will be adversely influenced [14]. The goal of this report is to identify the optimal nutrient dose for microbial community culture. Methane and carbon dioxide yield rate were the indicators used to assess the results of experiments.

## 2. Materials and methods

### 2.1 Materials

The coal and coalbed water samples used in this study were collected from Laohutai Fm 103^#^ coalbed located in Shenyang province of China (GPS coordinates 41.830256, 123.957639). The depth of the Fm 103^#^ gas-coal sample was 550 meters and the thickness of the coal seam was 6-18 meters. Laohutai Mine, in the east part of Fushun near Shenyang, Liaoning province, opened in 1901. The mine has 55.6 million tons of recoverable coal reserves until 2004[15]. The samples were collected by the State Key Laboratory of Coal Resources and Safety Mining. The field studies did not involve endangered or protected species, and only small sample quantities were used. And no specific permissions were required for these locations sample collection and research.

The coal samples were obtained from a newly exposed coal seam in the heading face by the channel sampling method following ASTM D 4596-86 with nitrogen protection. Two sample points were selected from the Fm 103^#^ coal seam. These samples were sealed in gas-tight steel canisters and set with gas-tight valves (manufactured by J&D Technologies) which enabled inert gas to flush the canister and protected with nitrogen immediately. The coal to be used for the microbe community source was crushed into small pieces (10-15 mm in diameter) at an aseptic bench (manufactured by JingXue Technologies). Nitrogen protection was used to maintain the coal in a continuous anaerobic environment. The sample was sealed in a sterilized gas desorption canister (manufactured by J&D Technologies) to desorb the absorbed gas until no gas desorbed in atmospheric pressure at 25 °C[16].

The formation of water samples from the Fm 103^#^ coalbed were collected in the same coal-bed in sterile glass bottles (manufactured by Fisher Scientific) which were filled to overflow to prevent oxygen ingress. Coalbed water (≤ 24 hours after collection) was autoclaved at 121 °C for 45 minutes and sealed in sterile glass bottles. Argon was infused to seal the top space of the bottles. The coalbed water samples were stored at 4 °C for no more than 3 days.

### 2.2 Culture

Five different nutrition culture media were tested in these experiments: MAC-1, MAC-2, MAC-3, MAC-4, MAC-5 [4,17] (MAC is the abbreviation of “the influence test of organic matrix and cations”). The final concentrations of the compounds (g/L) were as list in table 1, and 0.1 mL/L of resazurin was added as an oxygen indicator. Yeast extract was produced by Fisher BioReagents, and other chemicals were supplied by Acros Organics.

**Table 1.**
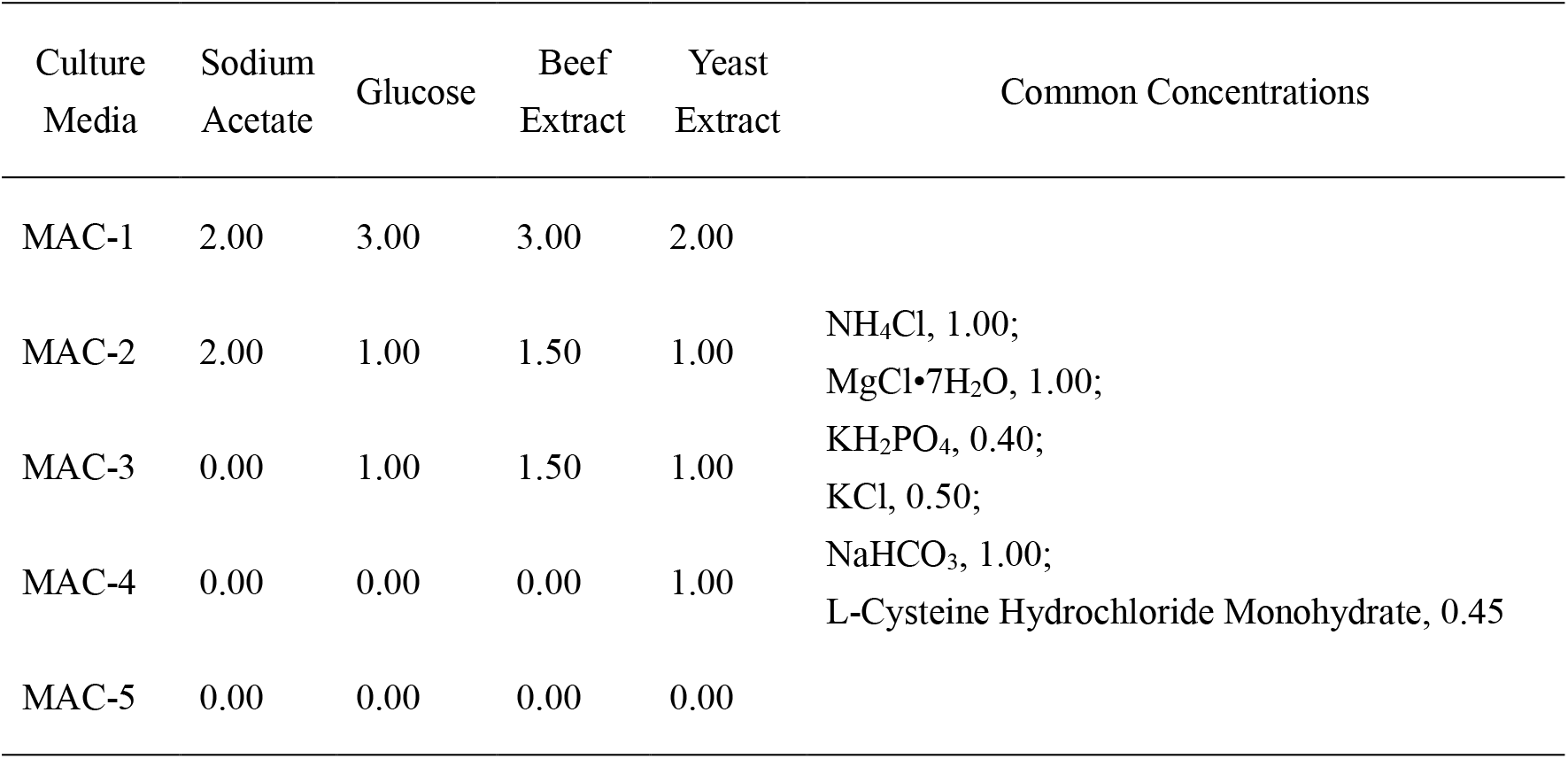
The MAC1-MAC5 nutrition culture media concentrations (g/L)

| Culture Media | Sodium Acetate | Glucose | Beef Extract | Yeast Extract | Common Concentrations |
| --- | --- | --- | --- | --- | --- |
| MAC-1 | 2.00 | 3.00 | 3.00 | 2.00 |  |
| MAC-2 | 2.00 | 1.00 | 1.50 | 1.00 | NH<sub>4</sub>Cl, 1.00;<br>MgCl•7H<sub>2</sub>O, 1.00; |
| MAC-3 | 0.00 | 1.00 | 1.50 | 1.00 | KH<sub>2</sub>PO<sub>4</sub>, 0.40;<br>KCl, 0.50; |
| MAC-4 | 0.00 | 0.00 | 0.00 | 1.00 | NaHCO<sub>3</sub>, 1.00;<br>L-Cysteine Hydrochloride Monohydrate, 0.45 |
| MAC-5 | 0.00 | 0.00 | 0.00 | 0.00 |  |

The distilled water was autoclaved at 121 °C for 45 minutes with dissolved oxygen removed. Nutrition medium was prepared in a 500 mL flask with distilled water. The medium was mixed using a magnetic stirrer for 2 hours at 60 °C at an aseptic bench and then combined with an equal volume of coalbed water (volume ratio 1:1) for another hour at room temperature. Nitrogen protection was used to maintain nutrition in a continuous anaerobic environment throughout the entire experiment. The final pH was maintained at 6.0 for all nutrition cultures. Control samples of 100 ml MAC-1, MAC-2, MAC-3, MAC-4, and MAC-5 were sealed in separate sterile glass bottles and stored at −40 °C. Argon was infused to seal the top space of bottles.

Anaerobic conditions were ensured in flasks using a gas-replacement method. This gas replacing process was monitored in real time with a carbon dioxide monitor system (manufactured by E2V). For each experiment 50.00±1 g of the coal sample and 500.00 mL of medium were used. Nitrogen was used to seal the upper space of the flask at the beginning of the experiment. The flasks were placed in an incubation shaker at 35 °C and were agitated at 80 rpm to maximize the coal-liquid mass transfer rates. Twelve parallel tests were designed for each nutrition group which were cultured for 40 days under identical conditions.

The control experiments were performed without coal sample, which named MAC-1*, MAC-2*, MAC-3*, MAC-4*, MAC-5*, to identify whether exogenous organic has the possibility to supply extra carbon to enhance the bio-methane yield. For each control experiment of each nutrition group 100 mL of cultured medium and 400 mL new medium was used. And cultured for 40 days in the same condition as above.

### 2.3 The cation orthogonal analysis

The cation orthogonal analysis was based on consumption of cation elements in bio-degradation processes, according to the L_16_(4^5^) orthogonal table which includes Na^+^, K^+^, Ca^2+^, Mg^2+^. The lowest ion level was based on the MAC-4 medium while the highest ion level was based upon that of the East China Sea ion concentration. A quantity of 50.00±1 g of Fm 103^#^ gas-coal sample was used in each experiment. The upper space of the flasks was sealed with nitrogen. The flasks were placed on an incubation shaker at 35 °C and agitated at 80 rpm to maximize coal-liquid mass transfer rates. Cation orthogonal experiments were of 40 days of culture.

### 2.4 Gas analysis

Gas samples were obtained with a 50-μL gas syringe. Methane and carbon dioxide analyses were performed using an Agilent 7890A gas chromatograph (manufactured by Agilent). The nitrogen (carrier gas) flow rate was set at 1.00 mL/min. The injection port was maintained at 150°C with the oven temperature set at 25°Cand the Thermal Conductivity Detector (TCD) at 200°C. Retention times for methane were 3.76 minutes and 5.0 minutes for carbon dioxide. Calibration standards consisted of 40% methane, 20% carbon dioxide, 10% hydrogen and 30% nitrogen, which were injected at atmospheric pressure to generate the calibration plot.

### 2.5 Nutrient Metabolism Analysis

Nutrient ion concentrations were analyzed using an HC-800 ionization analyzer (manufactured by Histrong Technologies). The main indicators included fluoride, chloride, nitrate and nitrite nitrogen, phosphate, sulfate, carbonate, bicarbonate, ammonia nitrogen, sodium, potassium, magnesium, calcium, pH, water hardness and total alkalinity.

## 3. Results

### 3.1 The nutrient abundance influence for bio-degradation of coal

Five different organic compound nutrient concentrations designated as MAC-1, MAC-2, MAC-3, MAC-4, and MAC-5 were assessed in this study. In these medium, Sodium Acetate is a collective medium, as carbon resource, for *methanogenic* normally [18]. Glucose is the medium for *hydrogen-producing acetogens*. Beef Extract is a mixture of peptides and amino acids, nucleotide fractions, organic acids, minerals, and some vitamins. It is often used to supply carbon and nitrogen sources. Yeast extract contains a mixture of amino acids, peptides, water soluble vitamins, and carbohydrates. And it is often used in culture media [19]. The rank order of organic concentrations were MAC-1 > MAC-2 > MAC-3 > MAC-4 > MAC-5.

Coalbed methanogenic groups are comprised of a variety of microbial types. With regard to *methanogens*, only a limited number of simple carbon compounds such as CO_2_ or acetate can serve as substrates. For the conversion of complex organic compounds to methane, fermentative and *acetogenic bacteria* are required. Thus they group an interactive methanogenic consortium [20]. In the process of gas-coal bio-degradation, the capacity for fermentative bacteria to hydrolyze and ferment the organic compounds of coal plays an important role, and CO_2_ is the main compound of gas productions. Beef extract in medium supplied the extra carbon and nitrogen for bacteria. For the *acetogenic bacteria* fermentation process, the long chain fatty acids and sugar degrade to form acetate, CO_2_ and H_2_ [18]. Glucose is the medium to enhance the *acetogenic bacteria* fermentation. And *methanogens* yield CH_4_ with CO_2_ and H_2_ or acetate. Sodium Acetate was introduced to raise the acetate supply for *methanogens* [21].

If consortia in a good banlance, most CO_2_ and H_2_ will be used to format CH_4_. And the concentration of CH_4_ should be high, meanwhile, the concentration of CO_2_ and H_2_ should keep in low. So the yield gas concentration with microbial fermentation process could reflect the bacteria balance conditions. The presence of sufficient organic nutrients in the medium could promote fermentative and *acetogenic bacteria* flourish and improve methanogen nutrient generation. However, if excessive amounts of organic nutrients are contained within the medium, which like beef extract and glucose, the high propagation of fermentative or *acetogenic bacteria* would break the microbial balance, and carbon dioxide yield rate could be enhanced, meanwhile, methanogenesis could be inhibited. This phenomenon showed in MAC-1, MAC-2 and MAC-3 culture experiments (Figure 1).

The results confirmed that MAC-4 was the most effective medium for enhancing the bio-methane generation rate. 1 g/L yeast extract provided the best concentration of organics amino acids, peptides, and water-soluble vitamins to optimize the microbial fermentation. The maximal methane concentrations of the MAC-4 culture group achieved 23.62% (Figure 1). It was 4 times higher than that of the MAC-2, MAC-3, MAC-5 groups on average. The MAC-4* control experiment and MAC-5 group verified that the anthracite is important to provide carbon for bio-methane yield (Figure 1).

**Figure 1.**
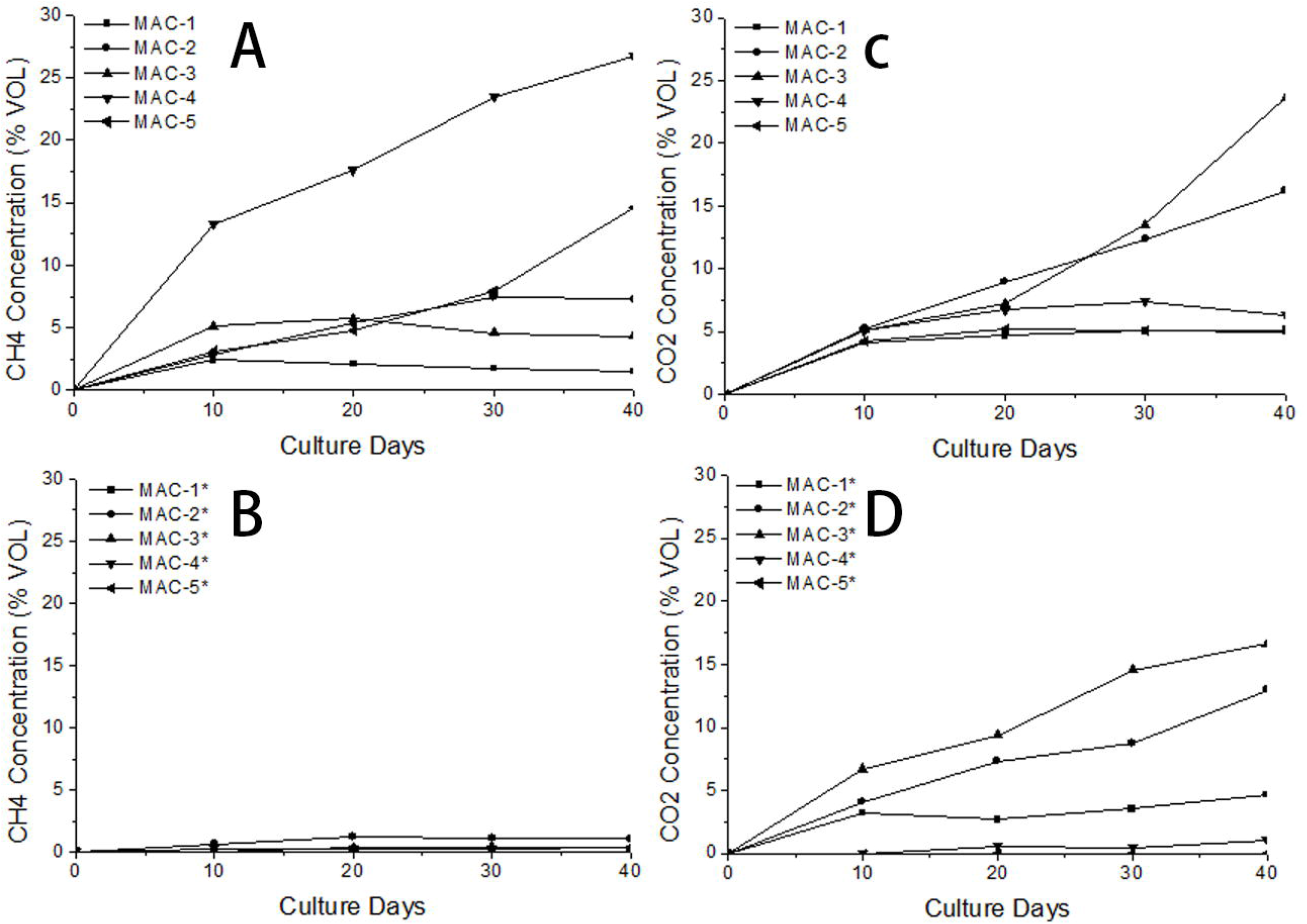
Changes in methane and carbon dioxide production as a function of media nutrition and time in culture. Figure A is the methane concentration changes with culture days. And figure B is the control experiment without coal sample supply in experiment. Figure C is the carbon dioxide concentration changes with culture days. And figure D is the control experiment without coal sample supply in experiment. Methane is the main factor to identify the *methanogens* activity. And carbon dioxide is an important factor to identify the fermentative and *acetogenic bacteria* activity. The methane yield rate would high, only if the fermentative and *acetogenic bacteria* activity in a limited condition. Organic material could enhance the fermentative and *acetogenic bacteria*. However, if the activity of fermentative and *acetogenic bacteria* is too high, it would inhibit the *methanogens*. The microbial group will in a good balance when achieved the highest methane yield and the best ratio of methane and carbon dioxide. Thus the MAC-4 medium culture group fit for the requirements.

The high carbon dioxide and low methane concentration data obtained from MAC-1 and MAC-2 media demonstrate that hydrolytic bacterial had enhanced with an abundance of organic substrate supplement, however, methane biosynthesis tended to inhibit in culture even with additional sodium acetate. In MAC-1 and MAC-2 experiments, the *acetogenic bacteria* inhibited also with sodium acetate influence in the medium.

The MAC-3 medium, which follows the same medium concentration of MAC-2 except sodium acetate, enhanced *acetogenic bacteria*, and inhibited the *methanogens*. After 40 days of culture, the average H_2_ volume concentration for the MAC-3 and MAC-3* group achieved 67.2% and 59.15% as indicated by gas analysis. This amount was 33-40 times greater than that of all other experimental groups. The exogenous organic material played an important role in carbon supply.

However, organic nutrition plays an important to supply the vitamin and other microelements besides organic nutrients. Without the vitamin and microelements besides organic nutrients supply, the consortia would grow at a slower rate, such as that observed for MAC-5.

### 3.2 The nutritional metabolism analysis of methanogenic consortium

Changes in MAC-4 ion concentrations were analyzed in the initial and final media. The data from this analysis revealed that the concentrations of sodium, potassium, ammonia nitrogen and magnesium were consumed microbially. In particular, 85.78% of the sodium was utilized with microbial fermentation. The medium pH changed from 6.65 to 7.32 over this period. In contrast, the concentrations of sulfate and bicarbonate increased 1.5-7 fold (Table 2). These data indicated that sodium, nitrogen, potassium, and magnesium are the key elements for methanogenic consortium fermentation. However, the sulfate should be maintained at a low level in the medium.

**Table 2.** The MAC-4 medium ion concentration changes with methanogenic consortium bio-degradation.

|  | pH | Fluoride | Chloride | Nitrate<br>Nitrogen | Nitrite<br>Nitrogen | Phosphate | Sulfate | Carbonate |
| --- | --- | --- | --- | --- | --- | --- | --- | --- |
| Initial | 6.65 | 3.70 | 81.58 | - | - | 44.67 | 195.80 | - |
| final | 7.32 | 5.18 | 95.90 | - | - | 38.60 | 478.40 | - |
|  | Bicarbonate | Sodium | Ammonia<br>Nitrogen | Potassium | Magnesium | Calcium | Water<br>Hardness | Total<br>Aalkalinity |
| Initial | 40.66 | 811.90 | 193.60 | 159.00 | 38.89 | 21.06 | 212.56 | 40.66 |
| final | 328.77 | 115.40 | 39.23 | 35.30 | 14.30 | 22.76 | 115.62 | 328.77 |

### 3.3 The cations orthogonal analysis

From the analysis of nutrient element concentrations, it is clear that sodium, ammonia, nitrogen, potassium, and magnesium play important roles in the methanogenic microbial metabolism. The orthogonal analysis experiment was designed to identify the influence of cations in bio-methane yield. The orthogonal module introduced 4 cations, (Na^+^, K^+^, Ca^2+^, Mg^2+^) as tested with 4 different concentrations. The lowest ion concentration was set by MAC-4 and the highest reference level for cation analysis was that of the East China Sea ion concentration. The L_16_(4^5^) orthogonal table was used as an orthogonal analysis module (Table 3).

**Table 3.**
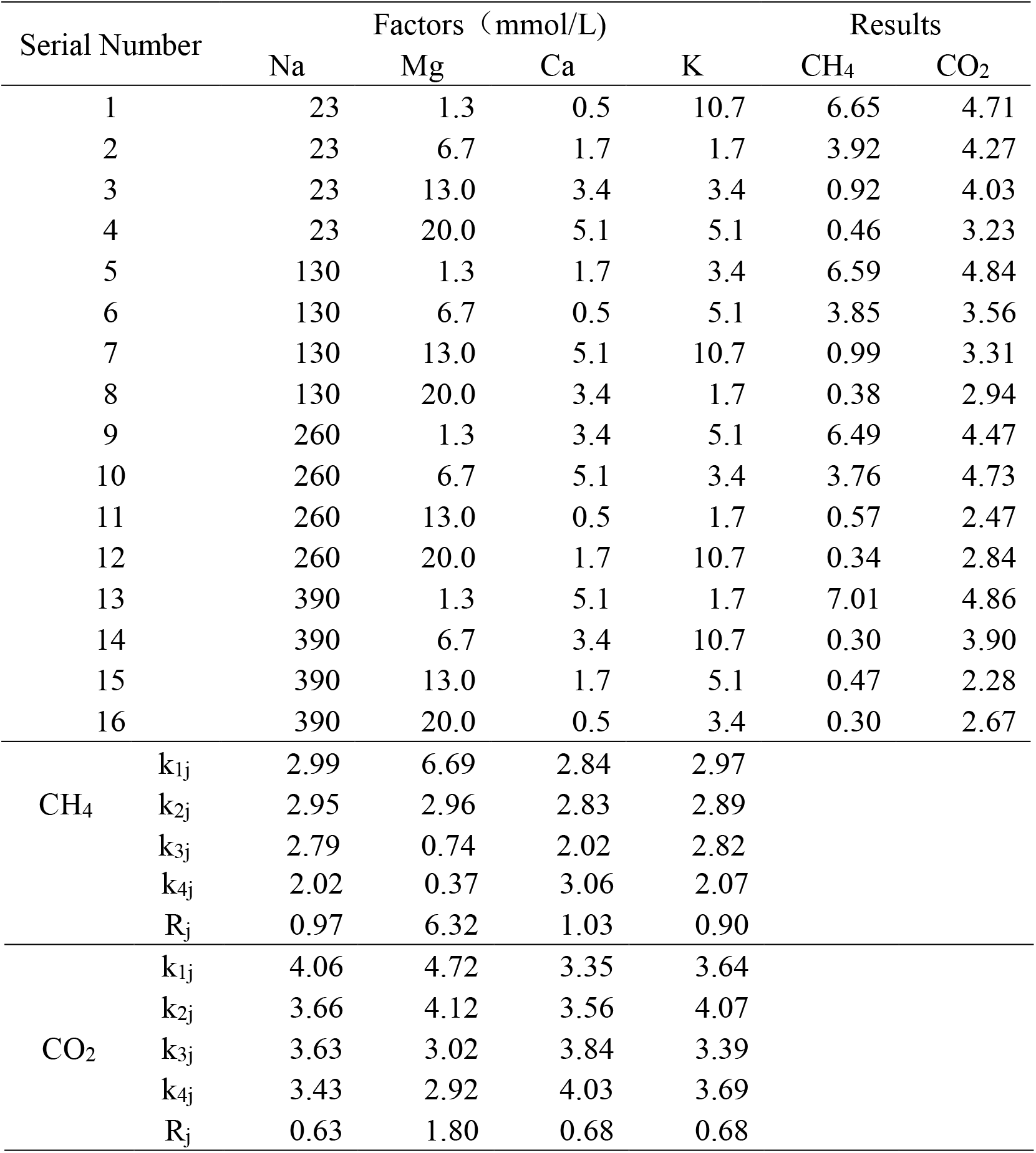
The cation orthogonal analysis table for CH_4_ and CO_2_. The results represent volume percent units for both gases. The table shows the analysis of CH_4_ and CO_2_ ranges.

Range analysis was used to indicate the affected order among factors. Range R_j_ was calculated with module 1.

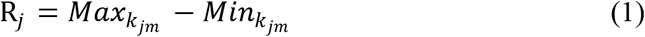

Where *k_jm_* is the average result of the *m* level *j* factor (module 2), (j, m =1, 2, 3, 4).

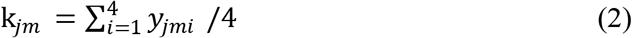

Where *y_jmi_* is the result of the *m* level *j* factor number i data, (j, m, i =1, 2, 3, 4).

Range analysis confirmed that for methanogenic activity, the best ion concentrations for enhancement CH_4_ yield were Na_1_, Mg_1_, Ca_4_, and K_1_, with the rank order of element effectiveness being Mg > Ca > Na > K. The most effective ion concentration for control CO_2_ generation were Na_4_, Mg_4_, Ca_1_, and K_3_, with its ran order being Mg > Ca = K > Na.

Notable differences exist regarding the influence of CH_4_ and CO_2_ on yield rates as a function of cation concentrations. The CH_4_ biosynthesis is substantially more sensitive to element content. Maximal Mg^2+^ concentrations to enhance methanogen metabolism processes were 1.3 mmol/L, while maximal cation concentrations for Ca^2+^, K^+^, Na^+^ were 5.1, 1.7 and 23 mmol/L, respectively.

Variance analysis has been used to analyze the Factor Significance (F) in this system. F was calculated with module 3.

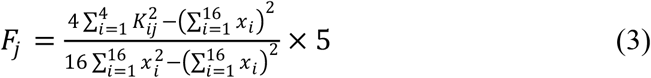

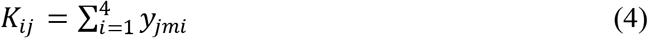

Where *x_i_* is the experiment results for every factor and level.

Based upon the F calculation, Mg^2+^ was the significant cation factor for CH_4_ and CO_2_ (Table 4). This ion is particularly sensitive in inhibiting CH_4_ metabolism processes largely for gas-coal methanogenic consortium.

**Table 4.** The variance analysis table for 4 factors with CH_4_ and CO_2_ data.

|  | Factor | Square of Deviance | Degree of Freedom | F Value | Significant |
| --- | --- | --- | --- | --- | --- |
| CH<sub>4</sub> | Na<sup>+</sup> | 2.465 | 3 | 0.892 |  |
|  | Mg<sup>2+</sup> | 100.905 | 3 | 36.533 | * |
|  | Ca<sup>2+</sup> | 2.486 | 3 | 0.900 |  |
|  | K<sup>+</sup> | 2.080 | 3 | 0.753 |  |
|  | Error | 2.762 | 3 |  |  |
| CO<sub>2</sub> | Na<sup>+</sup> | 0.842 | 3 | 8.592 |  |
|  | Mg<sup>2+</sup> | 9.120 | 3 | 93.061 | ** |
|  | Ca<sup>2+</sup> | 1.079 | 3 | 11.010 | * |
|  | K<sup>+</sup> | 0.954 | 3 | 9.735 |  |
|  | Error | 0.098 | 3 |  |  |

## 4. Discussion

Microbial bio-degradation of some organic compounds of coal represents a current technology available to enhance coal-bed methane. Microbial cooperation via anaerobic digestion processes results in the implementation of secondary biogenic methane generation and gas reservoirs enhancement in mining. Based on the experiments performed in this report, the following conclusions can be garnered: (1) The organic nutrient dose should be adjusted to methanogenic consortium requirements. Excessive or deficient organic nutrient support doses adversely affecting biomethane yield; (2) MAC-4 was the most effective medium in enhancing anaerobic digestion in Fushun gas-coal seam; (3) The sodium, nitrogen, potassium, and magnesium are the key elements for methanogenic consortium fermentation; (4) The cation rank order of influence Fushun methanogenic consortium metabolism was Mg > Ca > Na > K, and maximal cation concentrations of Mg^2+^, Ca^2+^, K^+^, Na^+^ were 1.3, 5.1, 1.7, and 23 mmol/L, respectively; (5) Mg^2+^ was a particularly sensitive factor, which could inhibit methanogenic bacteria.

## 5. Conclusion

### 5.1 The nutrient concentration for gas-coal methanogenic consortium culture

The biomethane generation from coal involves a complex interaction between environmental factors and biotic communities. Hydrolytic fermentative bacteria, syntrophic *acetogenic bacteria*, methanogenic bacteria, and many other bacteria comprise the biotic community. The environment includes not only the physical factors but also the coal, coalbed water, coal-bed gas and other complex formations which we call the environment.

The medium injected into coal seam enriching nutrients of the coal seam. The series of experiments of this report confirm that nutrient media require a strict concentration of control to be effective, especially for those involving organic materials. If the organic compounds are too rich, like that modeled with our formulations of MAC-1, MAC-2, and MAC-3, the excessive levels of nutrients will adversely affect the microbial community structure. High rates growth of hydrolytic fermentative bacteria or *acetogenic bacteria* could inhibit methane biosynthesis. In contrast, if lack organic nutrition, such as that modeled in MAC-5, the potential for enhancing methane biosynthesis is low and the yield proceeds at a slow rate. Concentrations of nutrition required should differ as a function of: (1) the methane biosynthesis type, such as carbon dioxide reduction or acetate fermentation; (2) coal maturity grade, such as gas-coal, flame coal, bituminous coal or lignite. The exact organic nutrition required for different coal ranks needs to be identified to maximize bio-methane yield rates.

### 5.2 Ion concentration for bio-methane yield enhancement

Organic biological degradation to methane is a microbial cooperation process. Fermentative bacteria initially hydrolyze complex organic compounds to acetate, longer chained fatty acids, carbon dioxide, hydrogen, NH_4_^+^, and HS^−^. Syntrophic hydrogen-producing (proton-reducing) *acetogenic bacteria* reduce intermediary metabolites to acetate, carbon dioxide, and hydrogen. Hydrogen-utilizing *acetogenic bacteria* demethoxylate low molecular weight ligneous compounds and ferment some hydroxylated aromatic compounds. Carbon dioxide reduction methanogenic bacteria are dependent on hydrogen, produced by other bacteria, to reduce carbon dioxide or bicarbonate to methane. And acetate fermentation methanogen yields methane via acetate bio-degradation.

Different nutrients are required for different microbial activity in the community. Results from the cation orthogonal experiments revealed that cation concentrations critically influence the metabolism process of the microbial. Except for the examination of sodium, potassium, magnesium, calcium which analyzed in the cations orthogonal analysis, the nitrogen, yeast extract, salinity and pH could have the potential to influence the biomethane synthesis. More researches to identify the effects of ions upon the coal seam biotic community are needed to reveal the microbial activity in gas-coal beds.

## 7. Acknowledgments

The authors’ acknowledge the contributions of the following companies for allowing access to coal samples and other information used in this paper: Laohutai Mining. We thank, Ke Li, Yumei Jia’s assistance in this work.

This work was supported by the State Key Laboratory of Coal Resources and Safe Mining Subject (grant number SKLCRSM17KFA08 and SKLCRSM19X012), the Fundamental Research Funds for Central Universities (grant number 2014QNB41).

## References

1. Lv Y, Tang D, Xu H, Luo H. Production characteristics and the key factors in high-rank coalbed methane fields: A case study on the Fanzhuang Block, Southern Qinshui Basin, China. Int J Coal Geol. 2012; doi:10.1016/j.coal.2012.03.009

2. Park SY, Liang Y. Biogenic methane production from coal: A review on recent research and development on microbially enhanced coalbed methane (MECBM). Fuel. 2016. doi:10.1016/j.fuel.2015.10.121

3. Scott AR. Improving Coal Gas Recovery with Microbially Enhanced Coalbed Methane. Coalbed Methane: Scientific, Environmental and Economic Evaluation. 1999. doi:10.1007/978-94-017-1062-6_7

4. Green MS, Flanegan KC, Gilcrease PC. Characterization of a methanogenic consortium enriched from a coalbed methane well in the Powder River Basin, U.S.A. Int J Coal Geol. 2008; doi:10.1016/j.coal.2008.05.001

5. Penner TJ, Foght JM, Budwill K. Microbial diversity of western Canadian subsurface coal beds and methanogenic coal enrichment cultures. Int J Coal Geol. 2010; doi:10.1016/j.coal.2010.02.002

6. Strąpoć D, Picardal FW, Turich C, Schaperdoth I, Macalady JL, Lipp JS, et al. Methane-producing microbial community in a coal bed of the Illinois Basin. Appl Environ Microbiol. 2008;74: 2424–2432. doi:10.1128/AEM.02341-07

7. Li D, Hendry P, Faiz M. A survey of the microbial populations in some Australian coalbed methane reservoirs. Int J Coal Geol. 2008; doi:10.1016/j.coal.2008.04.007

8. McIntosh J, Martini A, Petsch S, Huang R, Nüsslein K. Biogeochemistry of the Forest City Basin coalbed methane play. Int J Coal Geol. 2008; doi:10.1016/j.coal.2008.03.004

9. Hall RO, Tank JL, Baker MA, Rosi-Marshall EJ, Hotchkiss ER. Metabolism, Gas Exchange, and Carbon Spiraling in Rivers. Ecosystems. 2016; doi:10.1007/s10021-015-9918-1

10. Beckmann S, Luk AWS, Gutierrez-Zamora ML, Chong NHH, Thomas T, Lee M, et al. Long-term succession in a coal seam microbiome during in situ biostimulation of coalbed-methane generation. ISME Journal. 2018. doi:10.1038/s41396-018-0296-5

11. Chen T, Zheng H, Hamilton S, Rodrigues S, Golding SD, Rudolph V. Characterisation of bioavailability of Surat Basin Walloon coals for biogenic methane production using environmental microbial consortia. Int J Coal Geol. 2017; doi:10.1016/j.coal.2017.05.017

12. Xiao D, Wang E-Y, Peng S-P, Wu J-Y. Responses of coal anaerobic fermentation fractures development. Meitan Xuebao/Journal China Coal Soc. 2017;42. doi:10.13225/j.cnki.jccs.2016.1109

13. Meng J, Nie B, Zhao B, Ma Y. Study on law of raw coal seepage during loading process at different gas pressures. Int J Min Sci Technol. 2015; doi:10.1016/j.ijmst.2014.12.005

14. Galagan JE, Nusbaum C, Roy A, Endrizzi MG, Macdonald P, Fitzhugh W, et al. The genome of M. acetivorans reveals extensive metabolic and physiological diversity. Genome Res. 2002; doi:10.1101/gr.223902

15. Wang E, Chen P, Li Z, Shen R, Xu J, Zhu Y. Resistivity response in complete stress-strain process of loaded coal. Meitan Xuebao/Journal China Coal Soc. 2014;39. doi:10.13225/j.cnki.jccs.2013.1070

16. Ali Shah F, Mahmood Q, Maroof Shah M, Pervez A, Ahmad Asad S. Microbial ecology of anaerobic digesters: The key players of anaerobiosis. The Scientific World Journal. 2014. doi:10.1155/2014/183752

17. Parks DH, Chuvochina M, Waite DW, Rinke C, Skarshewski A, Chaumeil P-A, et al. A standardized bacterial taxonomy based on genome phylogeny substantially revises the tree of life. Nat Biotechnol. 2018; doi:10.1038/nbt.4229

18. Zinder SH. Physiological Ecology of Methanogens. Methanogenesis. 1993. doi:10.1007/978-1-4615-2391-8_4

19. Bellou S, Baeshen MN, Elazzazy AM, Aggeli D, Sayegh F, Aggelis G. Microalgal lipids biochemistry and biotechnological perspectives. Biotechnology Advances. 2014. doi:10.1016/j.biotechadv.2014.10.003

20. Gerritsen J, Fuentes S, Grievink W, van Niftrik L, Tindall BJ, Timmerman HM, et al. Characterization of Romboutsia ilealis gen. nov., sp. nov., isolated from the gastro-intestinal tract of a rat, and proposal for the reclassification of five closely related members of the genus Clostridium into the genera Romboutsia gen. nov., Intestinib. Int J Syst Evol Microbiol. 2014; doi:10.1099/ijs.0.059543-0

21. Kwietniewska E, Tys J. Process characteristics, inhibition factors and methane yields of anaerobic digestion process, with particular focus on microalgal biomass fermentation. Renew Sustain Energy Rev. 2014; doi:10.1016/j.rser.2014.03.041

